# BioLLM: A Standardized Framework for Integrating and Benchmarking Single-Cell Foundation Models

**DOI:** 10.1101/2024.11.22.624786

**Authors:** Ping Qiu, Qianqian Chen, Hua Qin, Shuangsang Fang, Yanlin Zhang, Tianyi Xia, Lei Cao, Yong Zhang, Xiaodong Fang, Yuxiang Li, Luni Hu

**Affiliations:** College of Life Sciences, University of Chinese Academy of Sciences, Beijing 100049, China; BGI Research, Beijing 102601, China; BGI Research, Wuhan 430074, China; BGI Research, Shenzhen 518083, China; DSA Thrust, The Hong Kong University of Science and Technology (Guangzhou)

## Abstract

The application and evaluation of single cell foundational models (scFMs) present significant challenges stemming from the heterogeneity of architectural frameworks and coding standards. To address these issues, we introduce BioLLM, a framework facilitating the integration and application of scFMs in single-cell RNA sequencing data analysis. BioLLM provides a universal interface, bridging diverse scFMs into a seamless ecosystem. By mitigating architectural disparities and coding conventions, it empowers researchers with streamlined access to scFMs. With standardized APIs and comprehensive documentation, BioLLM streamlines model switching and comparative analyses, while incorporating best practices for consistent model evaluation. Our comprehensive evaluation of scFMs revealed distinct strengths and limitations, highlighting scGPT’s robust performance across all tasks, both in zero-shot and fine-tuning scenarios. Geneformer and scFoundation also demonstrated strong capabilities in gene-level tasks, benefiting from effective pretraining strategies. In contrast, scBERT underperformed relative to other models, likely attributable to its considerably smaller parameter count and the limited size of the training dataset. Ultimately, BioLLM aims to empower the scientific community to leverage the full potential of foundational models, advancing our understanding of complex biological systems through enhanced single-cell analysis.

## Introduction

Single-cell RNA sequencing (scRNA-seq) has revolutionized molecular biology by enabling high-resolution transcriptome profiling, offering new insights into the complexity of biological systems^1, 2, 3, 4^. However, as vast amounts of single-cell data accumulate, effectively mining and extracting key features from these datasets pose significant challenges in biological research^5, 6, 7, 8, 9^. The advancement of deep learning, particularly through foundation models^10, 11, 12, 13, 14, 15^, presents substantial potential in addressing these challenges. Foundation models are AI systems trained on extensive and diverse datasets without reliance on human annotations, enabling them to capture complex patterns and adapt to a wide range of tasks with significantly less data compared to models trained from scratch^16, 17, 18^. Central to this innovation are Transformer^19^ architectures, exemplified by models such as BERT^20^ and GPT-4^21^, which have transformed the landscape of machine learning. The inherent flexibility and computational efficiency of Transformers make them exceptionally well-suited for the intricate task of mining single-cell data, thereby facilitating the extraction of meaningful biological insights from complex cellular landscapes^22, 23, 24, 25, 26^.

Leveraging the potential of foundation models to address the complexities of single-cell data analysis, several models, such as scBERT^10^, Geneformer^11^, scGPT^12^, and scFoundation^13^, have been developed to tackle specific challenges in this field (Table S1). However, these models demonstrate both commonalities and distinctions in their architectural design and pre-training strategies, accompanied by differences in dataset size and parameter count. For example, scBERT employs a bidirectional transformer trained by masked language modelling and incorporates gene2vec^27^ embeddings to represent gene identities^10^, while scGPT employs an autoregressive training strategy with Flash-Attention^28^ blocks and random gene identity embeddings, focusing on generating summaries for each cell^12^. Given these differences mentioned beforehand, the performance of each model can vary significantly across various downstream tasks, such as batch effect correction and cell type classification. Therefore, it is essential to evaluate these models systematically to determine which performs best in specific contexts.

Moreover, the varying levels of code accessibility and documentation across these single-cell foundation models (scFMs) create significant challenges for their unified implementation. While some models, such as Geneformer and scGPT, provide extensive documentation and well-structured open-source repositories that facilitate easy integration and customization, others may lack comprehensive guidelines or present less user-friendly implementations. This inconsistency can hinder researchers from effectively leveraging multiple models, as differing frameworks and coding standards lead to compatibility issues. Consequently, integrating these diverse models into a single analytical pipeline becomes challenging, limiting the ability to conduct comparative analyses across multiple tasks simultaneously. This lack of standardization not only affects the usability of the models but also complicates reproducibility and collaboration within the research community. To fully realize the potential of foundation models in single-cell RNA sequencing, it is essential to advocate for a standardized framework for model usage and integration. Such a framework would enable researchers to seamlessly call upon various models in a unified context, thereby enhancing the reproducibility of research findings and fostering greater collaboration.

Here, we introduce BioLLM, a standardized framework designed to facilitate the integration and utilization of foundation models in single-cell RNA sequencing analyses. BioLLM aims to provide a cohesive interface that allows researchers to easily access various scFMs, regardless of their underlying architectural differences or coding standards. By offering a set of standardized APIs and comprehensive documentation, BioLLM enables streamlined model switching and comparative analyses. Additionally, BioLLM incorporates best practices for model evaluation, ensuring that users can assess the performance of different models on specific tasks in a consistent manner. This framework not only simplifies the technical challenges associated with disparate models but also promotes collaboration by providing a common platform for researchers to share their findings and methodologies. Ultimately, BioLLM aspires to empower the scientific community to leverage the full potential of foundation models, driving forward advancements in the understanding of complex biological systems through single-cell analysis.

## Results

### BioLLM provides a unified framework for scalable scFM analysis

Single-cell foundation models represent a breakthrough in cellular heterogeneity analysis^29^, yet their widespread utilization faces three critical challenges: inconsistent preprocessing pipelines, heterogeneous model interfaces, and non-standardized evaluation metrics. To address these limitations, we developed BioLLM, a unified framework that standardizes the deployment of single-cell foundation models through three integrated modules **(Fig. 1)**. The first module implements a decision-tree based preprocessing interface that establishes rigorous quality control standards for input data **(Supplementary Fig. 1a)**. The BioTask Executor functions as the central analytical engine of the framework, implementing a systematic workflow that progresses through five stages: configuration parsing, model initialization, data preprocessing, dataloader construction, and task execution. This sophisticated pipeline facilitates both zero-shot inference via cell or gene embeddings and targeted model fine-tuning for specialized applications, including cell type annotation and drug response prediction. At the core of BioLLM lies its foundation model loader, which provides a unified interface for seamlessly integrating prominent single-cell foundation models such as scBERT, Geneformer, scFoundation, and scGPT **(Supplementary Fig. 1b)**. This standardized approach enables systematic deployment and comparative evaluation of multiple foundation models within a consistent analytical framework. The third module complements this architecture by implementing comprehensive performance metrics that assess three crucial aspects: embedding quality through silhouette scores, biological fidelity through gene regulatory network analysis, and prediction accuracy through standard classification metrics.

**Figure 1.**
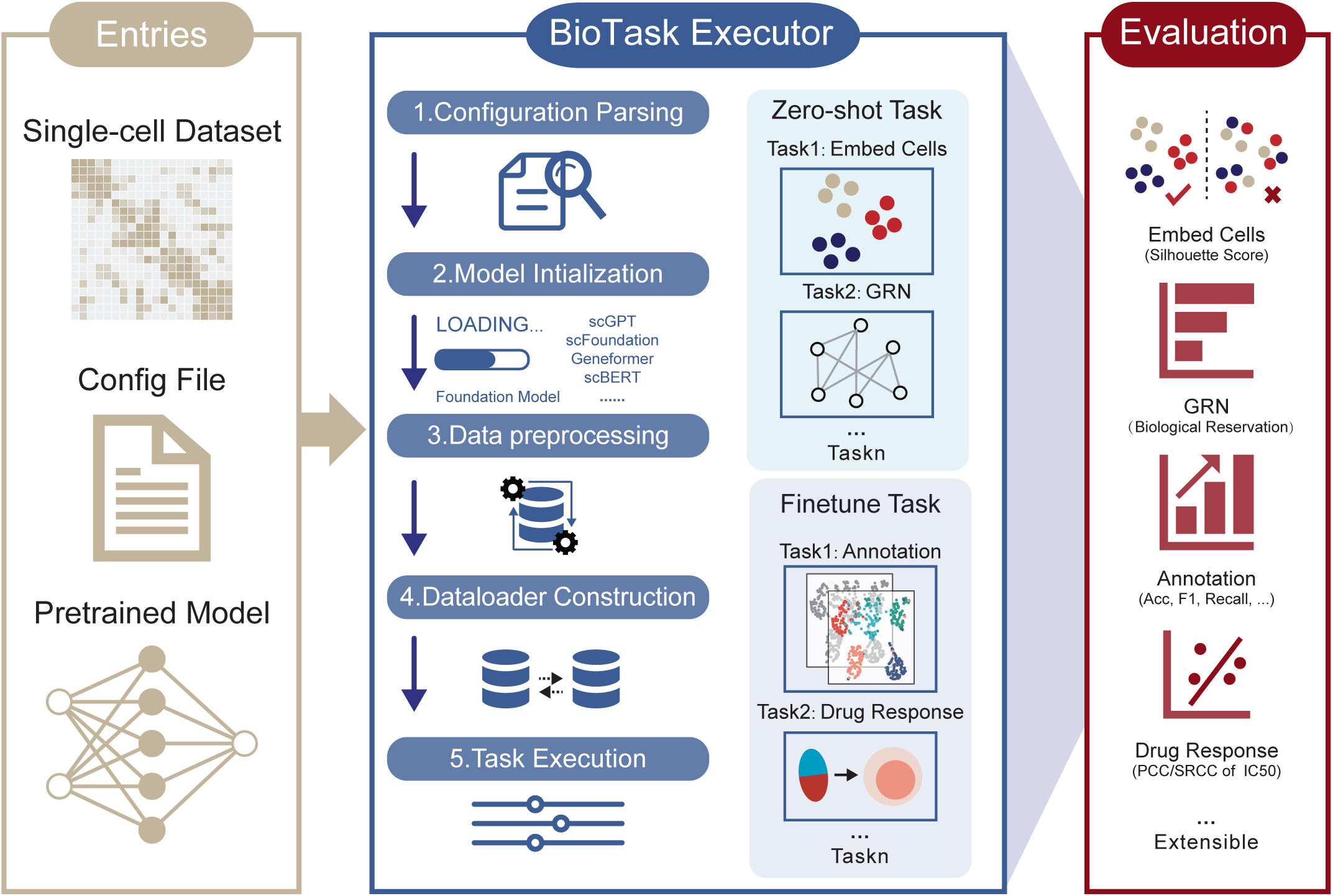
BioLLM Framework for Single-Cell Data Analysis. The BioLLM framework consists of three components: **Entries, BioTask Executor**, and **Evaluation**. Entries include the **input dataset, configuration file**, and **pretrained model**. The BioTask Executor processes tasks through five steps: **Configuration Parsing, Model Initialization, Data Preprocessing, Dataloader Construction**, and **Task Execution** (including zero-shot and fine-tuning tasks). Evaluation involves **cell embedding** (Average Silhouette Width, ASW), **GRN analysis** (Gene Ontology enrichment), **cell type annotation** (accuracy, precision, recall, macro F1), and **drug response prediction** (PCC, SRCC).

Through this integrated approach, BioLLM advances the field by providing a standardized, reproducible framework for large-scale single-cell data analysis across multiple foundation models, addressing a critical need in single-cell genomics research.

### BioLLM supports a comprehensive evaluation of cell representation capacity of scFMs

Single-cell foundational models leverage extensive training datasets to learn and generate cell embeddings, effectively transforming potentially noisy gene expression data into a biologically meaningful latent space^30^. We evaluated the performance of these models in zero-shot settings by assessing the quality of cell embeddings in both individual dataset and joint dataset contexts **(Supplementary Table 1)**, utilizing average silhouette width (ASW) as the evaluation metric. Our initial evaluations comprised four distinct individual datasets to confirm the biological relevance of the zero-shot cell embeddings. The results demonstrated that scGPT consistently outperformed other models **(Fig. 2a)**. UMAP visualizations further revealed that scGPT achieved superior separation of cell types compared to other foundational models **(Supplementary Fig. S2)**. This advantage can be attributed to scGPT’s capacity to capture complex cellular features, thereby enhancing separability. Its architecture is particularly proficient at preserving biologically relevant information, rendering it more effective for clustering tasks.

**Figure 2.**
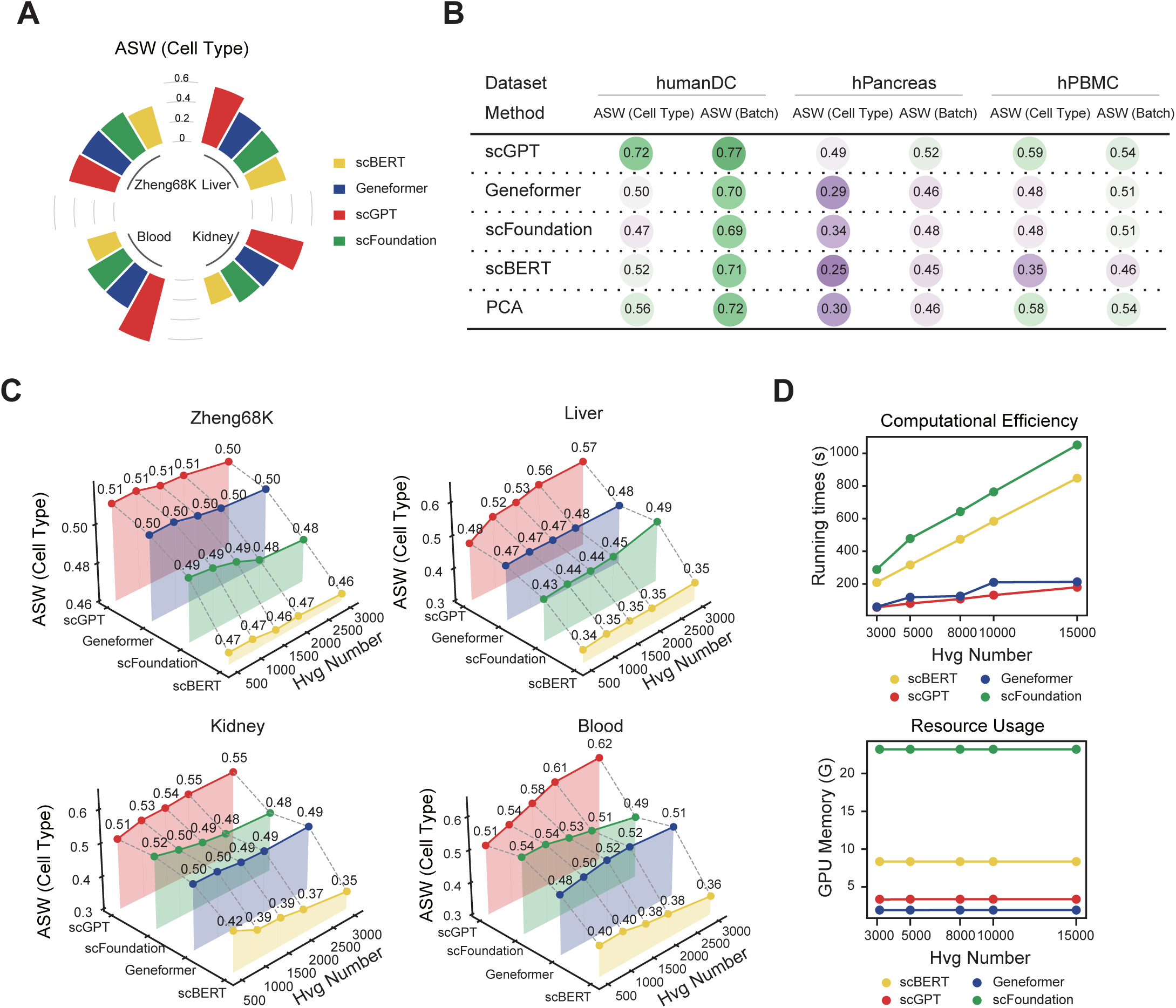
Evaluation of Cell Representation by Foundational Models. (A) Circular bar plot of ASW scores for scBERT, Geneformer, scGPT, and scFoundation across multiple datasets (Zheng68K, Liver, Kidney, Blood). (B) Summary table of ASW scores for cell type and batch correction across three datasets (humanDC, hPancreas, hPBMC). (C) 3D surface plots of ASW variation with the number of highly variable genes (HVGs) from 500 to 3000. (D) Line charts of running time (top) and GPU memory usage (bottom) for cell embedding with varying gene lengths. Evaluation metrics show scGPT’s superior cell type separation and computational efficiency.

Batch effects pose significant challenges in single-cell datasets, potentially compromising accurate interpretations. Consequently, a primary objective in single-cell analysis is to mitigate batch effects while preserving essential biological distinctions, thereby ensuring effective data integration^31, 32^. We evaluated the batch effect removal capabilities of single-cell foundational models in zero-shot cell embedding tasks using three joint datasets characterized by varying degrees of batch effects. Average Silhouette Width (ASW) scores, incorporating both cell type and batch information, were analyzed. Notably, scGPT outperformed the other models across both metrics, yielding superior results compared to PCA, while the other models performed worse than PCA **(Fig. 2b)**. UMAP visualizations demonstrated that while scGPT effectively integrated cells of the same type under consistent experimental conditions, it generally struggled to correct for batch effects across different technologies, whereas Geneformer and scFoundation distinguished certain cell types, but scBERT exhibited particularly poor performance **(Supplementary Fig. S3)**.

We further investigated the impact of varying gene input lengths on the cell embeddings generated by each foundation model **(Fig. 2c)**. The results indicated that as the input sequence length increased, scGPT embeddings became more accurate in representing true biological features, suggesting that longer input sequences enable scGPT to capture richer information, resulting in more accurate cell representations. In contrast, Geneformer and scFoundation exhibited a slight negative correlation between input length and embedding quality in some datasets, although the overall changes were minimal. Notably, scBERT’s performance declined as input sequence length increased across most datasets, potentially due to its difficulty in learning meaningful cell features, which led to more inconsistent embeddings. Additionally, we assessed the computational efficacy and resource usage associated with generating cell embeddings across the models **(Fig. 2d)**. Both scGPT and Geneformer demonstrated superior efficiency in terms of memory usage and computational time compared to scBERT and scFoundation, underscoring their practicality for large-scale analyses.

Overall, our evaluations demonstrate that BioLLM serves effectively as a comprehensive framework for assessing cell embeddings derived from various single-cell foundational models. While scGPT excels in generating biologically relevant embeddings and accurately distinguishing between cell types, it faces challenges in handling batch effects. Conversely, models like Geneformer and scFoundation demonstrate competitive performance, whereas scBERT shows notable deficiencies in this area.

### BioLLM facilitates in-depth analysis of gene regulatory networks across scFMs

Zero-shot gene-level evaluation is essential for benchmarking foundational models. BioLLM leverages gene-level embeddings to construct gene regulatory networks (GRNs), enhancing our understanding of gene interactions and regulatory mechanisms (**Fig. 3a**). By deriving gene embeddings from four distinct single-cell foundational models, BioLLM infers GRNs that facilitate the exploration of gene-gene interactions in a biologically relevant context. To further validate these findings, we plan to test the models on immune-related human datasets. The biological significance of these inferred GRNs is assessed by examining their capacity to delineate regulatory pathways and define functional relationships among genes. The results indicate that scGPT, scFoundation, and Geneformer exhibit a greater number of enriched pathways across all clustering resolutions compared to scBERT (**Fig. 3b**), particularly at lower resolutions for gene module identification. Focusing on core gene regulatory modules, the analysis of networks targeting HLA-DRA reveals that both scGPT and Geneformer effectively group HLA family genes, such as HLA-DRB5, HLA-DPA1, HLA-DQB1, HLA-DMB, HLA-DQA1, and HLA-DRB1, demonstrating a higher degree of interaction (**Fig. 3c**).

**Figure 3.**
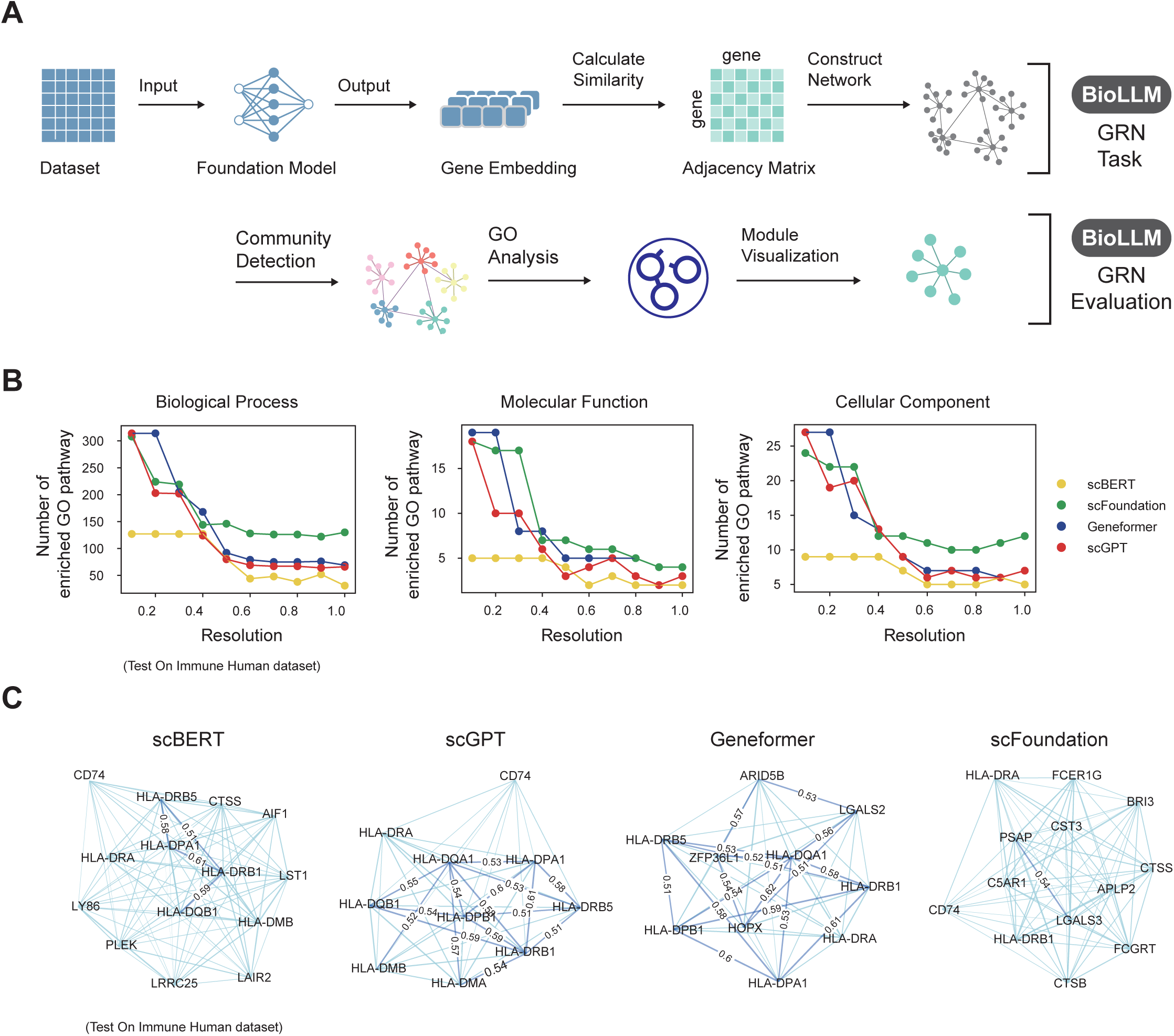
Gene Regulatory Network (GRN) Analysis. (A) Overview of GRN construction: gene embeddings are used to compute similarities and build the adjacency matrix, followed by community detection and Gene Ontology (GO) enrichment analysis. (B) Line plots of enriched GO pathways for each model at different resolution settings, indicating biological process, molecular function, and cellular component enrichment. (C) Network visualizations of HLA-DRA regulation for different foundational models, showing regulatory relationships among genes.

Collectively, these findings underscore the efficacy of BioLLM in utilizing gene-level embeddings to construct informative GRNs, thereby facilitating the identification of critical gene interactions and regulatory pathways. The enhanced performance of scGPT, scFoundation, and Geneformer in GRN construction emphasizes their potential to yield valuable biological insights into gene regulatory mechanism.

### BioLLM allows for comparative performance evaluation of cell annotation tasks among scFMs

As cell annotation is a crucial aspect of single-cell analysis, BioLLM incorporated this task to evaluate the performance of scFMs across 13 datasets from diverse tissues (Supplementary Table 1). The performance of these models was benchmarked against three established annotation methods: singleR^33^, celltypist^34^, and scANVI^35^. Four classification metrics were employed to rigorously assess model efficacy: accuracy, precision, recall, and macro-F1 score. The results indicated that scGPT outperformed all other foundational models, followed by Geneformer, scBERT, and scFoundation **(Fig. 4a and Supplementary Fig. 4)**. Compared to traditional annotation tools, scGPT consistently demonstrated superior performance, although Geneformer achieved a slightly lower F1 score than both celltypist and singleR. Additionally, the logistic regression-based celltypist outperformed both scBERT and scFoundation overall. Notably, in the context of rare cell type identification, scGPT exhibited greater capability than the other scFMs **(Fig. 4b, c)**. To simulate real-world scenarios, two datasets were allocated for cross-dataset evaluation, reflecting conditions where query data often suffers from batch effects relative to reference data **(Supplementary Fig. 5a)**. In this scenario, scGPT also exhibited superior annotation performance **(Supplementary Fig. 5b, c)**.

**Figure 4.**
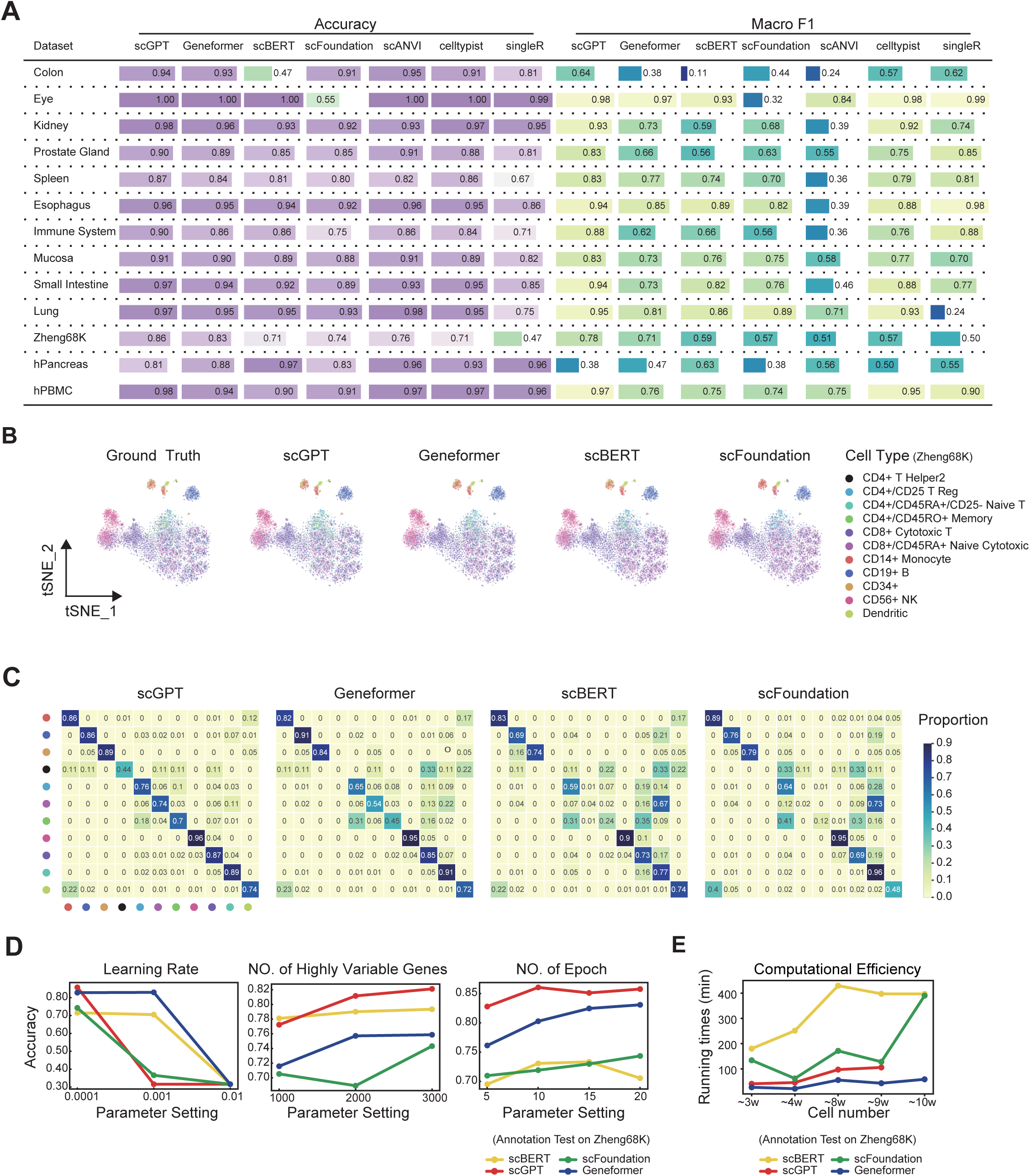
Cell Type Annotation Evaluation. (A) Overview of cell type annotation performance for four foundational models and three traditional methods across 13 datasets using accuracy and macro F1. (B) t-SNE plots of cells from the Zheng68K dataset, showing ground truth and model predictions. (C) Confusion matrices for Zheng68K dataset annotations by each model, comparing true and predicted cell types. (D) Line plots showing how learning rate, number of HVGs, and epochs influence annotation accuracy. (E) Computational efficiency comparison for different cell counts during model fine-tuning.

The impact of various hyperparameter settings on the performance of scFMs was also evaluated **(Fig. 4d)**. The results indicated that a lower learning rate and an increased number of training epochs generally improved model performance. Additionally, increasing the input gene sequence length positively influenced the annotation performance of scGPT, Geneformer, and scFoundation, while it had minimal effect on scBERT. Furthermore, we assessed the time required for the annotation task as the number of cells increased **(Fig. 4e)**. Geneformer achieved the shortest runtime, efficiently annotating 100,000 cells in under one hour, followed by scGPT. In contrast, scBERT required the longest time for the annotation process.

Taken together, the results highlight the effectiveness of BioLLM in utilizing scFMs for cell annotation tasks. Among these models, scGPT emerged as the preeminent model, demonstrating enhanced performance across multiple metrics and excelling in the identification of rare cell types.

### BioLLM enables seamless integration of scFMs and specific bioinformatics tools

To further investigate the capabilities of BioLLM, we aimed to determine whether integrating external bioinformatics tools during the fine-tuning of single-cell foundational models (scFMs) could enhance the framework’s applicability. In this study, we specifically replaced the transcriptomic feature extraction network in DeepCDR^36^ with four scFMs, utilizing their embeddings in subsequent DeepCDR network modules **(Fig. 5a)**. This integration was designed to predict the half-maximal inhibitory concentration (IC50) values of various drugs across multiple cell line datasets.

**Figure 5.**
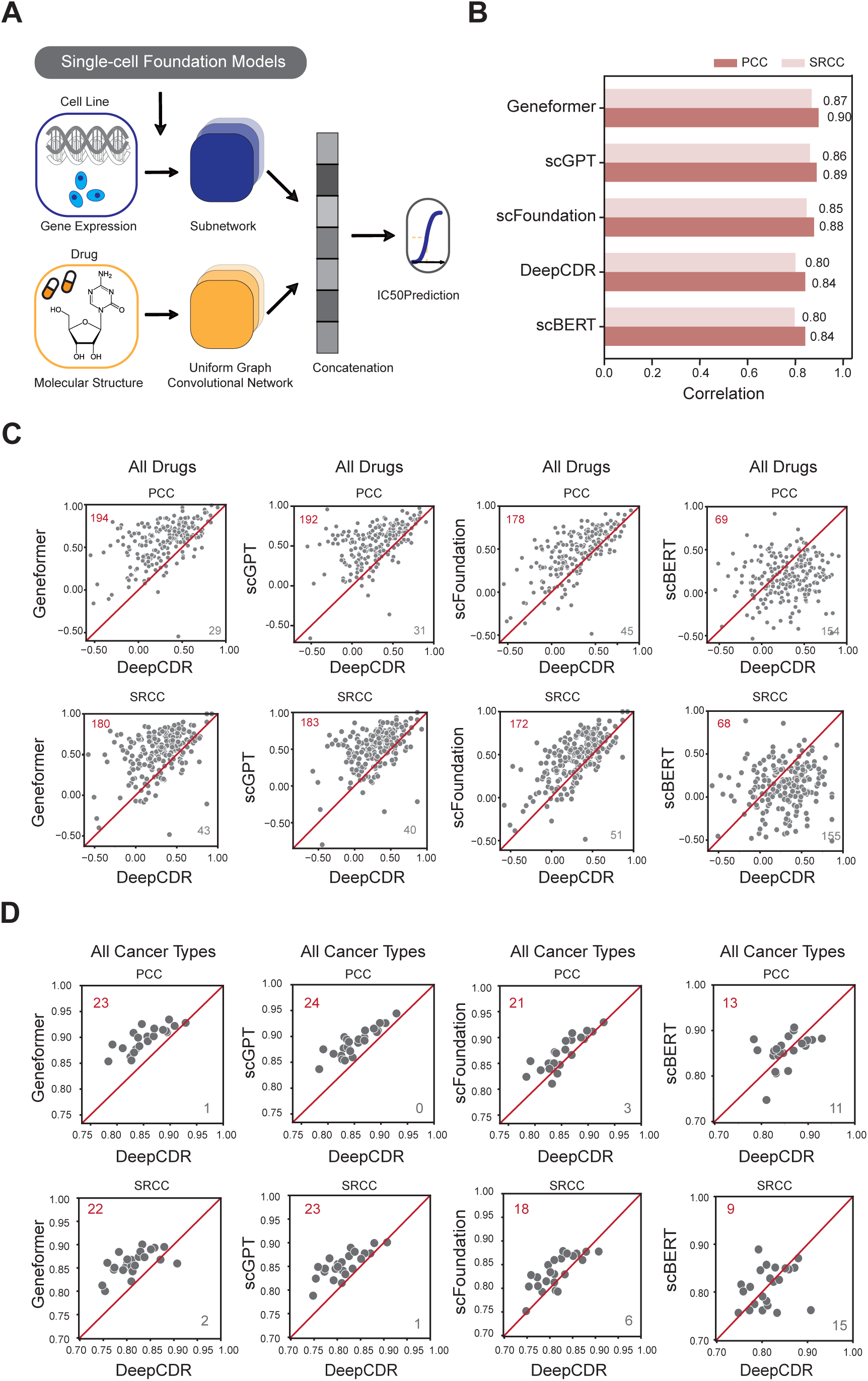
Cancer Drug Response (CDR) Prediction Using Foundational Models. (A) Schematic of drug response prediction workflow combining gene expression, drug molecular structure, and foundational models. (B) Bar plot of PCC and SRCC for foundational models compared to DeepCDR. (C) Scatter plots comparing PCC for all drugs between DeepCDR and foundational models. (D) Scatter plots comparing PCC and SRCC for all cancer types, highlighting improved accuracy for foundational models like Geneformer and scGPT.

We assessed the performance of these scFMs in comparison to the traditional DeepCDR model across various drugs and cell lines, using the Pearson correlation coefficient (PCC) and Spearman’s rank correlation coefficient (SRCC) as key performance metrics **(Fig. 5b and 5c)**. The results indicated that replacing the gene expression feature extraction in DeepCDR with scFMs generally led to improved performance, except for scBERT, which did not exhibit any significant enhancement. Notably, Geneformer and scGPT achieved the highest performance, producing comparable results, followed by scFoundation. Across all cancer types examined, embeddings from Geneformer and scGPT consistently yielded higher PCC and SRCC values **(Fig. 5d)**.

In conclusion, BioLLM facilitates the seamless integration of scFMs with external bioinformatics tools, thereby enhancing biologically relevant discoveries. The evaluation results underscore the effectiveness of scFMs in improved predictive performance for cancer drug response.

## Discussion

In this study, we introduced BioLLM, an integrative tool that consolidates multiple scFMs into a cohesive framework, thereby enhancing the analysis of single-cell RNA sequencing (scRNA-seq) data. BioLLM facilitates the streamlined selection and application of various models, specifically scBERT, Geneformer, scGPT, and scFoundation, enabling researchers to efficiently navigate the complexities inherent in single-cell genomics. By supporting both zero-shot learning and fine-tuning tasks, BioLLM not only optimizes research workflows but also ensures high-quality, reproducible outcomes in downstream analyses. This is particularly critical given the urgent need for standardized methodologies to manage the increasing volume and complexity of single-cell data.

Our comprehensive evaluation of the foundational models revealed distinct strengths and limitations that warrant further discussion. Among all benchmarked scFMs, scGPT demonstrated robust performance across all downstream tasks, excelling in both zero-shot and fine-tuning scenarios. This suggests that its generative pretraining is particularly effective for synthesizing biological insights from complex datasets. However, its ability to mitigate batch effects was found to be suboptimal, likely due to insufficient incorporation of batch-related information during pretraining. This limitation underscores the critical need to integrate batch-specific variability into model training to enhance performance in practical applications^37^.

Notably, Geneformer and scFoundation also exhibited commendable capabilities in gene-level tasks. Geneformer’s effective pretraining strategy, which focuses on the ordering of gene sequences and the prediction of gene IDs, significantly enhances its understanding of gene-gene interactions. This targeted approach appears to confer advantages in tasks that require nuanced knowledge of molecular relationships^38^. In contrast, scBERT exhibited significant underperformance relative to other scFMs, potentially due to its pretraining strategy for encoding full-length gene sequences, combined with limited parameters and insufficient training data, which impede their ability to accurately represent both cellular and gene-level relationships. These findings highlight the necessity for a critical reevaluation of the architectures employed in these models and suggest that refinements in pretraining methodologies could substantially improve their efficacy.

These findings highlight the necessity for ongoing refinement in the design and training of scFMs to overcome the limitations identified in our evaluations. Future research should focus on enhancing model specificity and generalizability, particularly across diverse datasets and biological contexts, to ensure that these models can be applied effectively in a variety of experimental scenarios. Additionally, exploring potential synergies between models could yield more comprehensive insights into biological systems at the single-cell level. Such integrative approaches may ultimately lead to the development of hybrid models that capitalize on the unique strengths of individual foundational models, thereby advancing the field of single-cell genomics^39^.

In summary, BioLLM establishes a standardized framework for integrating scFMs and facilitates biological exploration. This cohesive architecture not only streamlines model selection and application but also promotes consistency and reproducibility in single-cell analysis. By addressing the complexities of single-cell genomics, BioLLM empowers researchers to derive meaningful insights and fosters advancements in our understanding of biological systems at the single-cell level.

## Methods

### BioLLM Framework Design and Implementation

The BioLLM framework is structured around three core components: foundational architecture, task management, and model loading interfaces. This design emphasizes modularity, flexibility, and extensibility, facilitating the integration of diverse foundational models and a wide range of downstream tasks for single-cell data analysis.

#### Framework Architecture

BioLLM establishes a standardized infrastructure for managing configurations, loading models, and executing analytical tasks. This architecture promotes seamless interactions between model and task modules, enabling independent operation while ensuring consistent interoperability. Such a feature is critical in the rapidly evolving landscape of single-cell genomics, where adaptability is paramount.

#### Configuration Management

The framework includes a unified configuration management module that allows users to specify parameters via a configuration file or an API. This includes model selection, data preprocessing specifications, and task types (e.g., fine-tuning or zero-shot inference). The configuration manager efficiently parses these parameters and supplies the necessary inputs for model loading and task execution, thereby simplifying the setup process and enhancing user experience while minimizing errors.

#### Modular Task and Model Separation

To meet diverse analytical requirements, BioLLM implements a clear separation between task management and model loading. The task management module is responsible for data preprocessing, training, and inference, while the model module is dedicated to initializing and managing models independently. This organization not only enhances code structure but also allows users to customize and extend functionalities with ease.

#### LoadLlm Model Management Class

The LoadLlm class serves as the primary interface for loading and managing pre-trained foundational models within the BioLLM framework. Its key features include interface definition, compatibility checks, as well as data handling and embedding generation.

#### Interface Definition

LoadLlm defines essential interface methods such as load_pretrain_model(), load_data(), and get_embedding(). These methods are implemented in specific model subclasses, accommodating architectural variations across foundational models.

#### Data Handling and Embedding Generation

The class standardizes input data preprocessing to guarantee compatibility across models. A unified embedding generation process ensures that cell and gene embeddings produced by various models are appropriate for subsequent analyses, enhancing the reliability of downstream tasks.

#### BioTask Class for Task Management

The BioTask class is central to managing downstream analytical tasks specific to single-cell data and incorporates several key functionalities.

#### Configuration Parsing

BioTask loads and parses the configuration file to identify essential parameters, including task types (fine-tuning or zero-shot), model selection, and preprocessing requirements. Based on these parameters, BioTask dynamically selects and initializes the appropriate LoadLlm subclass corresponding to the specified foundational model.

#### Compatibility Checks

During model loading, LoadLlm performs rigorous compatibility checks on model parameters and task specifications, ensuring that the selected model is suitable for the intended task type— whether for fine-tuning or zero-shot inference—thus minimizing potential runtime issues.

#### Data Preprocessing and Dataloader Creation

BioTask processes input data into the AnnData format, widely utilized in single-cell analyses, facilitating organized access to observations (cells) and variables (genes). The class standardizes raw input data to meet model requirements and constructs a dataloader to manage large-scale datasets efficiently during execution.

#### Task Execution

The execution logic within the BioTask class is tailored to the analytical task at hand. For fine-tuning tasks, BioTask loads the pre-trained model and adjusts its parameters to optimize performance on the target dataset. In contrast, for zero-shot tasks, it leverages the pre-trained model to directly extract cell and gene features, generating embeddings suitable for subsequent analyses without retraining.

#### Extensibility and Modular Design

BioLLM is designed to facilitate the rapid integration of new tasks and models, ensuring adaptability to emerging research needs.

#### Adding New Task Types

Developers can extend BioLLM’s functionality by defining new analytical tasks within the BioTask class. This is accomplished by implementing specific task execution methods, allowing for the seamless incorporation of additional task types without modifying the core framework logic.

#### Integrating New Models

The framework supports the introduction of new foundational models through subclassing the LoadLlm class and implementing required interface methods. This modular design ensures compatibility with existing BioLLM task structures, streamlining the model integration process and encouraging innovation in modeling techniques.

#### Documentation and User Guide

Comprehensive documentation accompanies the BioLLM framework, providing users with detailed guidance on configuring and executing tasks. This includes step-by-step instructions for model integration, task execution, and extending the framework to accommodate new analytical tasks. This resource enhances user experience, supporting both novice and experienced users in effectively utilizing BioLLM for advanced single-cell data analysis.

### Evaluation of Downstream Tasks

#### Cell Embedding

The input dataset underwent preprocessing tailored to the specific requirements of each foundational model. For Geneformer and scGPT, which support input sequence lengths of 2048 and 1200, respectively, we selected 3000 highly variable genes as input features. In contrast, the other two foundational models utilized full-length gene sequences without feature selection. The preprocessing steps adhered to each model’s pretraining conditions to ensure optimal performance.

Specifically, scBERT, scFoundation, and scGPT required a log1p transformation of the gene expression data, while Geneformer utilized raw counts without normalization. For the generation of cell embeddings, BioLLM supports three methods for scBERT: CLS, mean, and sum. The CLS method utilizes the CLS token embedding as the cell representation, while mean and sum pooling methods apply mean and sum operations over the token embeddings produced by the model’s encoder.

For scGPT, the cell embeddings are derived from the CLS token embedding from the original model. In the case of Geneformer, embeddings are extracted using the model’s native function, which computes the mean of non-padded token embeddings. For scFoundation, the final cell embedding is generated through max pooling applied to the token embeddings.

This comprehensive approach to preprocessing and embedding generation ensures that each model’s strengths are leveraged, facilitating accurate representation of cellular data for downstream analysis.

#### Gene Regulatory Network Analysis

The Gene Regulatory Network (GRN) analysis was conducted using the Immune_ALL_human dataset as the input, processed through foundational models integrated with the BioLLM framework. The input dataset was preprocessed based on quality criteria as follows: 1) genes that have at least 3 counts were kept; 2) the raw data was subset to 1200 highly variable genes. Once preprocessed, the dataset was passed through the foundational model, which generated gene embeddings as output. Following the embedding generation, Euclidean distance was computed to construct an adjacency matrix. The distance quantified the relationships between each gene, and the adjacency matrix was developed to represent these interactions. Each element within the matrix indicated the degree of similarity between gene pairs, thus highlighting potential regulatory relationships. This adjacency matrix then served as the foundation for constructing the gene regulatory network, wherein nodes represented genes and edges depicted their inferred regulatory connections.

#### Cell Type Annotation

Cell type annotation was conducted in both intra-dataset and inter-dataset scenarios to assess the performance of the foundational models. Each dataset was divided into training and testing sets at an 8:2 ratio, ensuring a robust evaluation of model performance. The training set was further subdivided into training and validation subsets, also using an 8:2 split.

During the preprocessing phase, all datasets retained their complete gene expression profiles without applying any selection criteria for highly variable genes. This approach aimed to provide a comprehensive input for model training, facilitating the evaluation of each model’s capacity to leverage the full range of gene expression data.

Model training involved optimizing two critical hyperparameters: the number of training epochs and the learning rate (lr). A uniform epoch count of 20 was applied across all models to standardize the training duration. The learning rates were set according to the default values recommended for each foundational model: 0.0001 for both scGPT and scFoundation, 0.001 for scBERT, and 0.00005 for Geneformer.

To further investigate the impact of hyperparameter settings on cell annotation performance, the **Zheng68k** dataset was employed. This dataset allowed for a detailed exploration of how variations in epoch numbers, learning rates, and the selection of variable genes influenced the efficacy of the cell annotation models, providing insights into the optimal configuration for accurate cell type classification.

#### Drug Response

To evaluate and validate drug sensitivity predictions across the four single-cell foundational models, we utilized two datasets from DeepCDR: the Genomics of Drug Sensitivity in Cancer (GDSC), which provides IC50 values for each drug-cell pair, and the Cancer Cell Line Encyclopedia (CCLE), containing comprehensive gene expression, mutation, and methylation data for various cancer cell lines.

The embedding module of DeepCDR comprises two primary networks. The first network is a Graph Convolutional Network (GCN) tasked with feature extraction from the drug’s feature matrix and adjacency matrix. Concurrently, three distinct sub-networks are employed to encode the gene expression, methylation, and mutation data of the cell lines, effectively capturing the latent biological characteristics of the cells. The encoded features from both the drug and the cell line are then concatenated and processed through a Convolutional Neural Network (CNN) to predict the IC50 values.

In the integration of single-cell foundational models with DeepCDR, we focused on leveraging gene expression data from the CCLE dataset. Prior to inputting this gene expression data into the sub-networks of DeepCDR, we employed the selected single-cell foundational model to extract cell-specific features. These features were subsequently incorporated into the sub-networks, allowing for enhanced processing by the CNN to predict the IC50 values.

Model performance was rigorously assessed by calculating both the Pearson Correlation Coefficient (PCC) and Spearman’s Rank Correlation Coefficient (SRCC) between the predicted and actual IC50 values. These statistical measures provided a robust evaluation of the prediction accuracy, facilitating a comprehensive comparison of the models’ effectiveness in drug response assessment.

### Evaluation Metrics

#### Cell Embedding Metrics

To evaluate the effectiveness of cell embeddings, we assessed the separation of cell types and the presence of batch effects in the embedding space. The Average Silhouette Width (ASW) was employed as the primary metric, which quantifies the relationship between within-cluster and between-cluster distances. We computed ASW scores for both cell types and batch effects using a variant provided by scBI.

For cell types, the ASW score is normalized between 0 and 1, where a score of 0 indicates tight within-cluster cohesion, 0.5 suggests overlapping clusters, and 1 signifies well-separated clusters. Higher

ASW values indicate better cluster separation and overall performance. The ASW for cell types is calculated as follows:

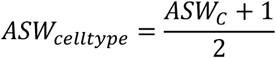

where *C* represents the set of all cell identity labels.

In the context of batch effects, ASW is computed based on batch labels, where a score of 0 denotes perfect mixing, and scores deviating from 0 indicate varying degrees of batch effects. To ensure that higher scores reflect better batch mixing, we scale the scores by subtracting them from 1, resulting in a range from 0 to 1. A score of 0 indicates complete batch separation, while a score of 1 reflects ideal batch mixing and integration.

#### Gene Regulatory Network Metrics

To assess the performance of the foundational models in gene regulatory network (GRN) analysis, we utilized the Immune_ALL_human dataset. The dataset was preprocessed according to the specific requirements of each model to ensure optimal embedding generation. Gene embeddings were then extracted to construct a neighborhood graph illustrating the connectivity among individual cells.

The Leiden algorithm was employed to cluster cells based on this connectivity, with the resolution parameter varied systematically from 0.1 to 1.0 in increments of 0.1. This approach allowed for an exploration of clustering at various levels of granularity, resulting in distinct subgroups that provided insights into cellular heterogeneity.

Subsequently, we filtered out clusters with more than 25 genes, and performed Gene Ontology (GO) enrichment analysis on the genes within each of these cluster, evaluating biological processes, molecular functions, and cellular components associated with these genes. The final result of GO enrichment only included categories with adjusted P values less than 0.01 This analysis offered valuable insights into the functional characteristics of the identified subgroups.

Additionally, we selected HLA-DRA as the target gene for visualization of the gene regulatory network. The visualization elucidated the regulatory relationships and interactions involving HLA-DRA, enhancing our understanding of its role in the context of immune response regulation.

#### Cell Type Annotation Metrics

Model performance in cell type annotation was assessed by comparing predicted results with true labels using four key metrics: accuracy, precision, recall, and macro F1 score. The macro F1 score, which accounts for class imbalances, is calculated by first determining the F1 score for each class. The F1 score is defined as the harmonic mean of precision and recall:

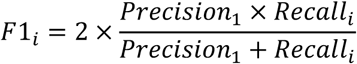

where *F*1_*i*_is the F1 score for class *i*, and*Precision*_*i*_ and*Recall*_*i*_ represent the precision and recall for class *i*. The macro F1 score is then computed by averaging the F1 scores across all *n* classes:

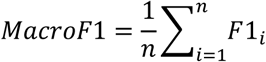

#### Drug Response Metrics

For the evaluation of drug sensitivity predictions, we utilized preprocessed data from DeepCDR. The primary metrics employed were the Pearson Correlation Coefficient (PCC) and Spearman’s Rank Correlation Coefficient (SRCC). These metrics assess the correlation between predicted and true IC50 values for each cell line across the foundational models, providing a robust measure of prediction accuracy. The PCC is calculated as follows:

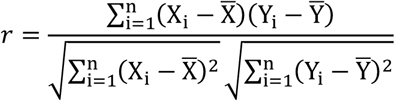

where *X*_*i*_ and *Y*_*i*_ refer to predicted IC50 value and the actual IC50 value for the i-th drug-cell pair separately. 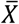 and 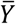 represent the mean value of all *X*_*i*_ and *Y*_*i*_.

The SRCC is calculated as follows:

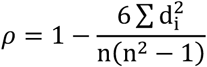

where *d*_*i*_ is the difference in ranks between the predicted IC50 value and actual IC50 value for the i-th drug-cell pair. n is the number of drug-cell pairs.

## Data availability

All data used in the research were collected from published sources, with detailed descriptions provided in Supplementary Table 2. For the evaluation of cell embedding, the Blood, Liver, and Kidney datasets, as well as a dataset accessible from the GEO database (GSE80171), were used. The datasets used for the cell type annotation task consist of the Spleen, Small Intestine, Prostate Gland, Immune System, Eye, Esophagus, Kidney, Colon, Lung, Zheng68K, 3’ dataset, 5’ dataset and four pancreas datasets. For the gene regulatory network task, the Immune_all_human was utilized. All datasets mentioned above can be downloaded from the links provided in Supplementary Table 2. The scFM-associated model files are available at http://doi.org/10.5281/zenodo.14189969.

## Code availability

BioLLM is packaged, and distributed as an open-source, publicly available repository at https://github.com/BGIResearch/BioLLM.

## Acknowledgement

We acknowledge the Stomics Cloud platform (https://cloud.stomics.tech/) for providing GPU computational resources. We thank the colleagues in our research group for inspiring discussion and their contributions.

## Author contributions

L.H. conceptualized the study. P.Q., and L.H. were responsible for the framework design and tool implementation. Q.C., and H.Q. performed data analysis and model evaluation. S.F., Y.L, T.X. and L.C. contributed key ideas and advice. P.Q., Q.C and L.N.H wrote the manuscript. L.H, Y.L, X.F, and Y.Z. supervised the study.

---

**Supplementary Figure 1.**
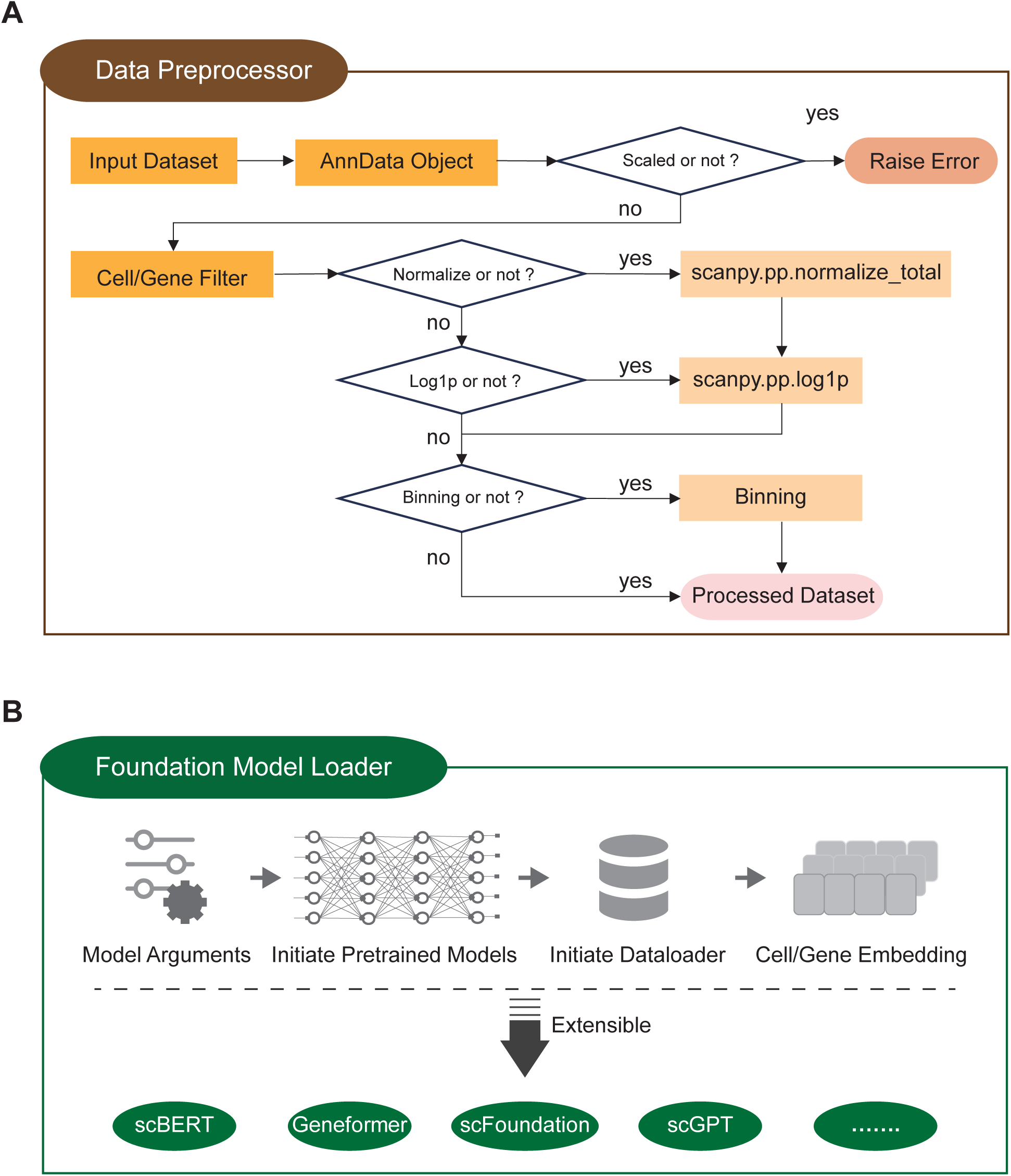
BioLLM Modules Overview. (A) **Data Preprocessor**: Data preparation workflow including AnnData conversion, scaling, normalization, log transformation, and binning for compatibility with models. (B) **Foundation Model Loader**: Process for loading pretrained models, initializing dataloaders, and generating embeddings, designed to support additional models.

**Supplementary Figure 2.**
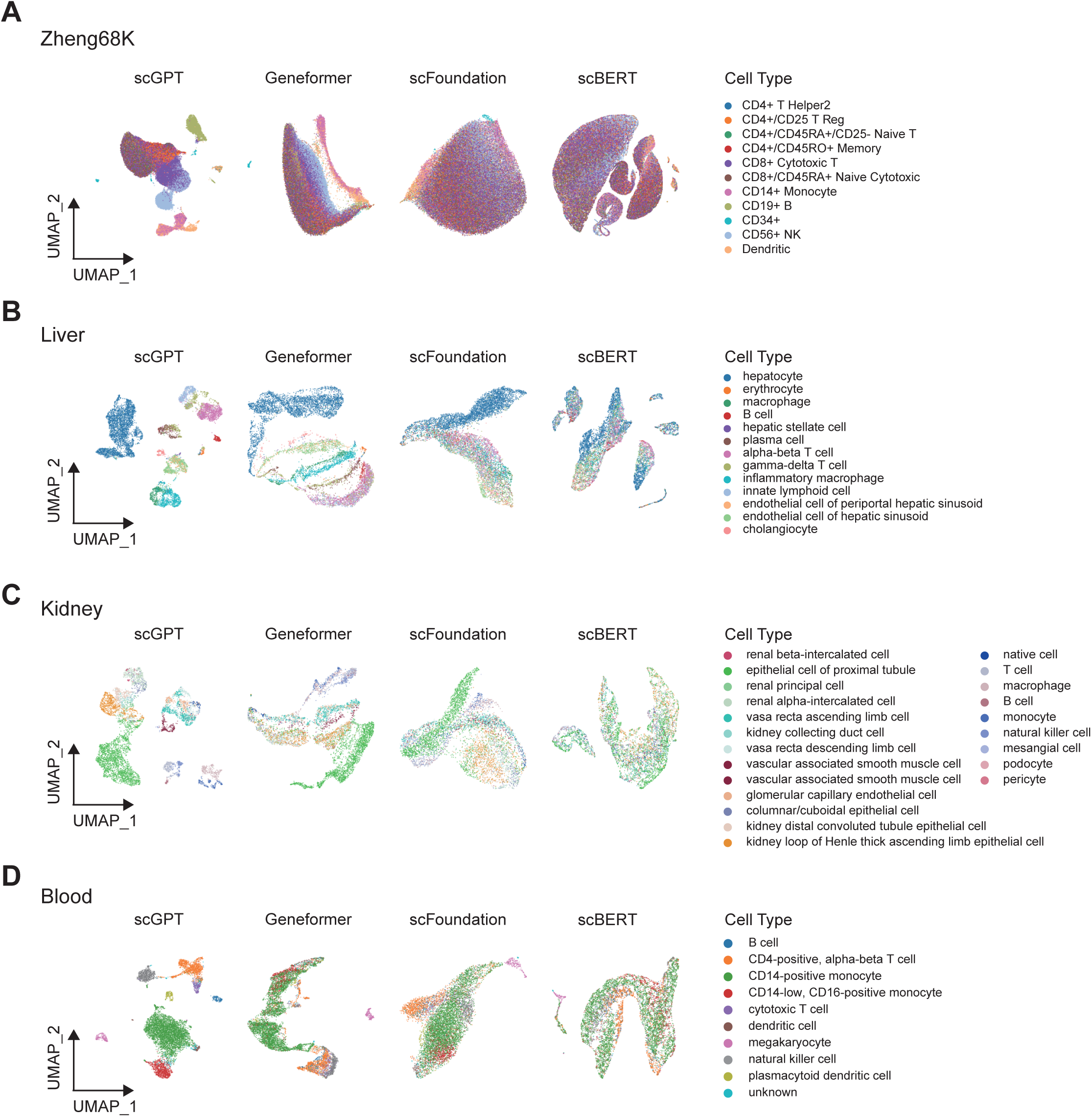
UMAP Visualization of Cell Embeddings. (A-D) **UMAP plots** of cell embeddings generated by scGPT, Geneformer, scFoundation, and scBERT for four datasets: (A) Zheng68K, (B) Liver, (C) Kidney, and (D) Blood.

**Supplementary Figure 3.**
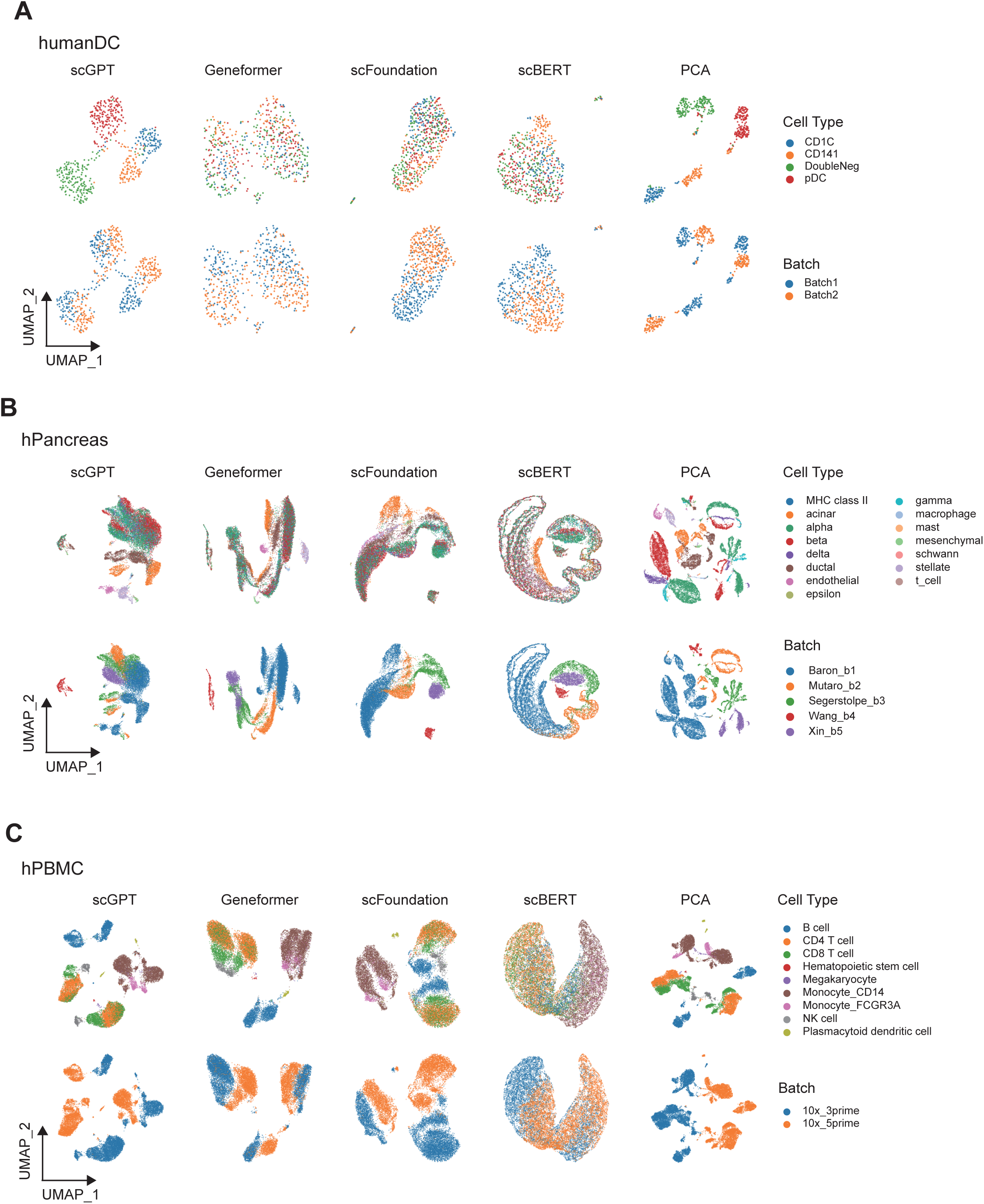
UMAP Plots Colored by Cell Types and Batches. UMAP visualizations for embeddings from (A) humanDC, (B) hPancreas, and (C) hPBMC datasets, colored by cell type and batch.

**Supplementary Figure 4.**
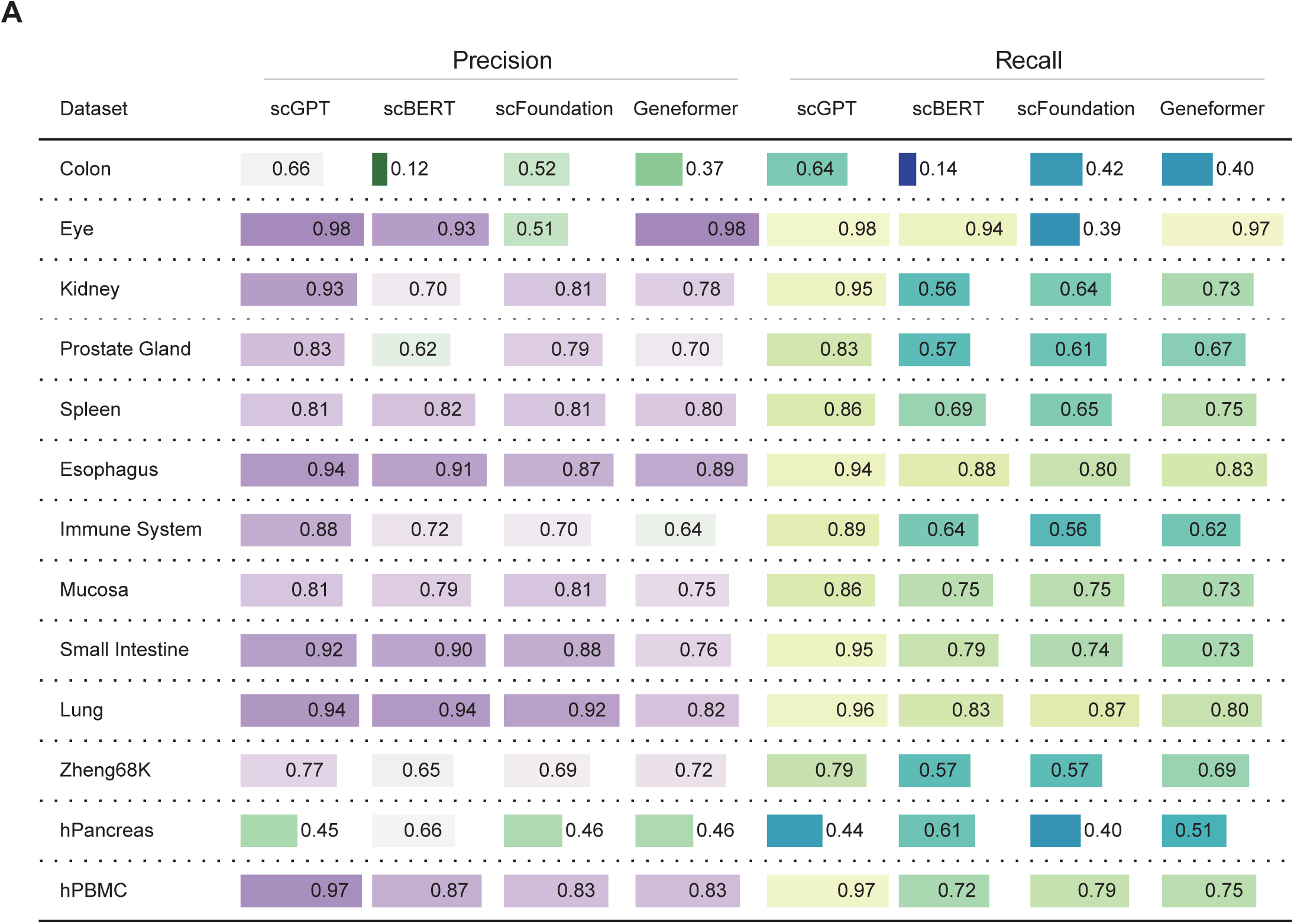
Cell Type Annotation Performance. Precision and recall metrics for foundational models in cell type annotation across 13 datasets.

**Supplementary Figure 5.**
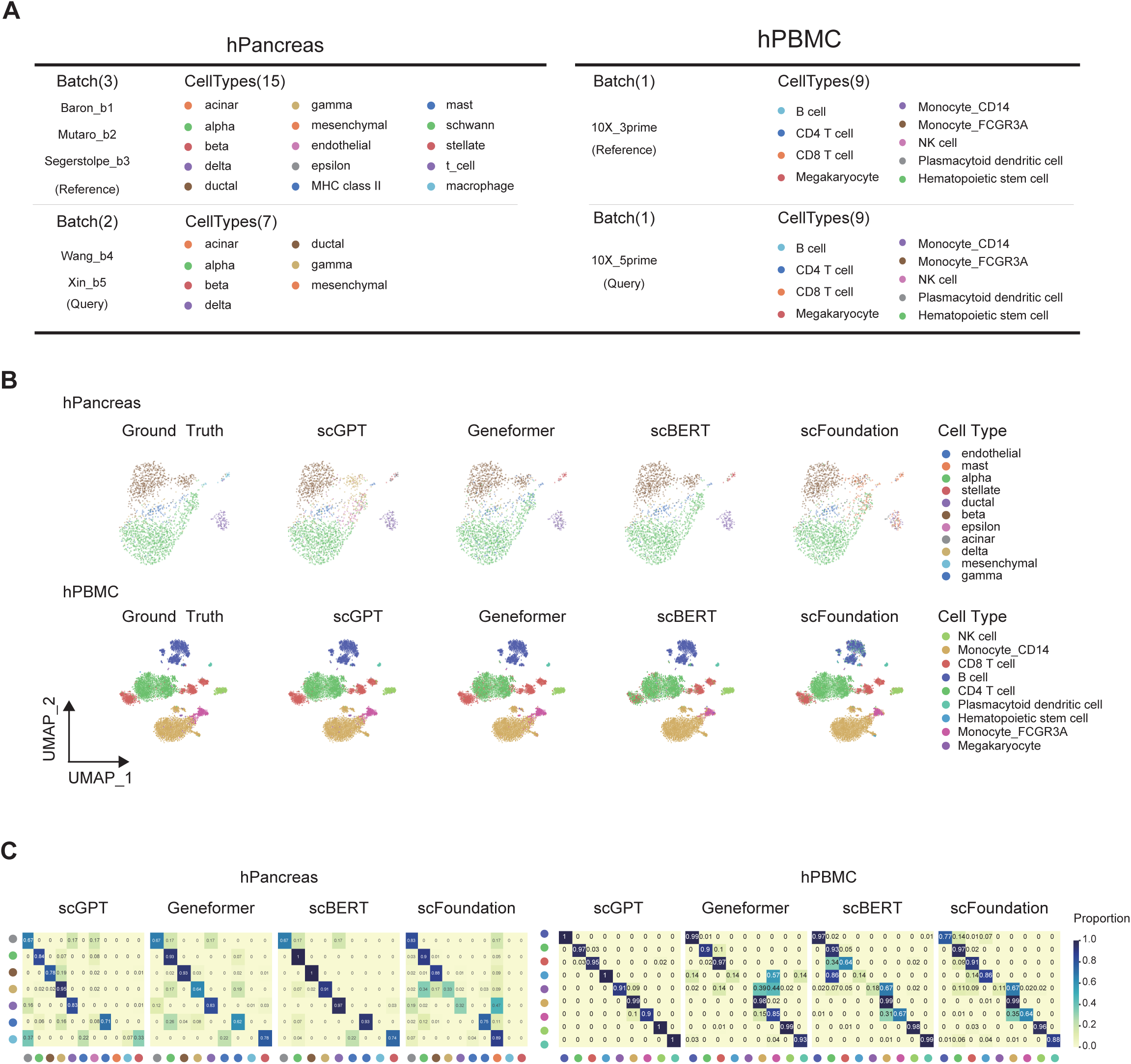
Cell Type Annotation Evaluation with Batch Splitting. (A) Batch-based data splitting for hPancreas and hPBMC. (B) t-SNE plots of cells from hPancreas and hPBMC with ground truth and model predictions. (C) Confusion matrices for hPancreas and hPBMC annotations by different models.

**Supplementary Figure 6. Drug Response Correlation Analysis.**

(A) PCC and (B) SRCC scatter plots comparing DeepCDR and foundational models for all drugs in the test set.

